# Transformed reactivation of latent working memory enables hierarchical language processing

**DOI:** 10.64898/2026.05.02.722174

**Authors:** Jiaqi Li, Yali Pan, Hyojin Park, Peter Hagoort, Huan Luo, Ole Jensen

**Affiliations:** Oxford Centre for Human Brain Activity, Oxford Centre for Integrative Neuroimaging, Department of Psychiatry, University of Oxford, Oxford, United Kingdom; Department of Experimental Psychology, University of Oxford, Oxford, United Kingdom; Centre for Human Brain Health, School of Psychology, University of Birmingham, Birmingham, United Kingdom; Max Planck Institute for Psycholinguistics, Nijmegen, The Netherlands; Donders Institute for Brain, Cognition and Behaviour, Radboud University, Nijmegen, The Netherlands; School of Psychological and Cognitive Sciences, Peking University, Beijing, China; PKU-IDG/McGovern Institute for Brain Research, Peking University, Beijing, China; Key Laboratory of Machine Perception (Ministry of Education), Peking University, Beijing, China

**Author notes:** **Correspondence:** (J.L.), (O.J.).

**Keywords:** language comprehension, working memory, activity-silent, reactivation, syntactic processing, prefrontal cortex, MEG, multivariate analysis

## Abstract

Human language comprehension requires tracking words across intervening material to construct grammatical structures, yet two fundamental questions remain unresolved: whether maintenance of earlier words relies on sustained neural activity or activity-silent working memory mechanisms, and whether memory retrieval during integration engages domain-general or language-selective networks. Using magnetoencephalography with time-resolved decoding, we tracked main-clause subjects as participants heard sentences with embedded clauses (e.g., “The dog, who chases the cat, jumps over the mud.”). The subject representation (“dog”) decayed to baseline during the embedded clause but reactivated after the main-clause verb (“jumps”), with transformed rather than reinstated neural codes. Critically, reactivation emerged first in right dorsolateral prefrontal cortex before engaging frontotemporal language regions, and reactivation strength was modulated by syntactic structure and predicted comprehension accuracy. These findings demonstrate activity-silent working memory maintenance, structure-dependent retrieval, and cooperative function of domain-general and language-selective networks during hierarchical language processing.

## Introduction

The human capacity for language depends on the ability to construct hierarchical structures that link grammatically related elements across intervening material ^1,2^. Resolving such dependencies requires keeping earlier words in memory while processing subsequent material, and later retrieved at appropriate points to enable flexible integration ^3,4^. However, how memory supports dynamic integration during sentence comprehension at the neural level remains largely unexplored ^5,6^. Long-distance dependencies, such as the subject-verb dependency between “the reporter” and“admitted” in “The reporter, who the senator attacked, admitted the error.”, provide a natural test case for investigating the neural implementation of working memory (WM) in sentence processing.

Previous studies on hierarchical structure building during sentence processing have yielded important insights into the timing and localisation of syntactic operations. Research in this area has identified event-related potential components, such as the Syntactic Positive Shift / P600, in response to syntactic violations ^7–10^, neural oscillations associated with structural processing ^11–13^, and demonstrated that syntactic operations can be decoded from neural activity ^14–16^. Yet these approaches do not directly reveal how content-specific linguistic representations are maintained across embedded structure and subsequently retrieved for integration.

How linguistic content is maintained in WM during sentence processing remains unresolved, but research in visual and auditory WM has identified two competing mechanisms about how the brain stores information over time. Persistent-activity theory proposes that WM representations are sustained through ongoing neural firing, maintaining information in an active state ^17–21^. In contrast, the activity-silent theory suggests that information can be stored in short-term synaptic plasticity without elevated firing to reduce interference between multiple items ^22–24^. These accounts make distinct predictions for hierarchical sentence processing. Persistent-activity models predict sustained neural representation of earlier linguistic elements throughout the embedded clause, whereas activity-silent theory predicts a loss of decoding accuracy followed by reactivation at integration. Although extensive research has revealed how linguistic information at various levels is encoded during comprehension and retrieved during production ^14,16,25,26^, few studies have investigated how word-level representations are maintained and integrated across long-distance dependencies and no study has directly test persistent-activity against activity-silent mechanisms during hierarchical sentence processing.

Another central unresolved question is whether WM during sentence comprehension is implemented within language-selective cortex alone, or whether it also engages domain-general WM regions, which support cognitive control and goal-directed behvior across diverse cognitive tasks ^27,28^. Converging evidence from neuroimaging and patient studies suggests that language-selective regions, particularly the left inferior frontal gyrus (LIFG) and temporal cortex, robustly respond to linguistic input across multiple levels of processing, including phonological, lexical, and syntactic analysis ^29–32^. However, the role of domain-general regions, including dorsolateral prefrontal cortex (DLPFC), in language comprehension remains highly debated ^33–35^. On the one hand, behavioural and neuroimaging evidence suggests that domain-general executive resources support core aspects of linguistic interpretation, including lexical retrieval, syntactic operation, and semantic integration ^30,36–39^. On the other hand, some researchers have questioned the extent to which the domain-general WM network support language comprehension, arguing against a role for this network in maintaining linguistic representations in WM ^25,33,34^. Here we address this question by investigating the engagement of domain-general regions during WM retrieval in long-distance dependency integration, and by examining the interaction between domain-general and language-selective networks.

To investigate the dynamic neural mechanisms by which WM supports sentence comprehension, we combined magnetoencephalography (MEG) with multivariate pattern analysis (MVPA) ^40,41^. MEG provides millisecond-level temporal resolution, enabling us to track moment-by-moment changes in neural representations as participants listen to normal-speed natural sentences. This fine-grained temporal precision is essential for capturing the rapid dynamics of memory retrieval at the precise moment when a long-distance dependency is resolved. Additionally, MEG offers sufficient spatial resolution to localise the brain regions involved in WM encoding and reactivation. Crucially, MVPA can detect word-specific information encoded in distributed neural activity patterns, even when univariate measures of overall activation appear unchanged ^42,43^. Furthermore, temporal generalisation of MVPA allow us to adjudicate between accounts that posit reactivation of the original encoding pattern at integration versus those that propose a transformed representation ^44^. Together, MEG analysed using MVPA provide the spatiotemporal precision and representational sensitivity needed to characterise both the neural substrates and the nature of WM processes during hierarchical sentence processing.

We recorded MEG from 34 native English speakers listening to spoken sentences containing embedded relative clauses (e.g.,“The dog, who chases the cat, jumps over the mud.”). Using time-resolved MVPA at both sensor and source levels, we tracked the neural representation of the main-clause subject (“dog”) from initial encoding, through the embedded clause (“who chases the cat”), to the point of subject-verb integration (“jumps”). We tested two key predictions. First, persistent-activity models predict sustained decoding of the subject throughout the embedded clause, whereas activity-silent models predict a transient loss of decoding accuracy followed by reactivation at integration. Second, if retrieval is implemented within language-selective cortex, reactivation should be confined to frontotemporal regions; if domain-general circuits contribute, reactivation should additionally engage DLPFC. Finally, we compared subject-relative and object-relative clauses to test whether the syntactic structure modulates WM reactivation and whether such modulation predicts comprehension accuracy.

## Results

### Behavioural performance in sentence-picture matching task

Participants (N=34, 20 females, aged 21.0 ± 3.0, mean ± SD) listened to English sentences and judged whether a subsequent probe picture matched the sentence meaning (Fig. 1A). Each sentence consisted of a main clause with a subject-verb-prepositional phrase structure, with either a subject-relative (SR) or object-relative (OR) clause embedded between the main-clause subject and verb (e.g., SR: “The dog, who chases the cat, jumps over the mud”; OR: “The dog, who the cat chases, jumps over the mud”). The main-clause subject was drawn from a set of four animals (dog, cat, fox, goat), enabling multivariate decoding of subject identity across trials. The embedded-clause noun was selected from the remaining three animals, and the main-clause verb was either “jumps over” or “steps into”, yielding 48 unique sentences. All the sentence constituents were counterbalanced across conditions (see full details in Methods and Supplementary Table 1).

**Figure 1.**
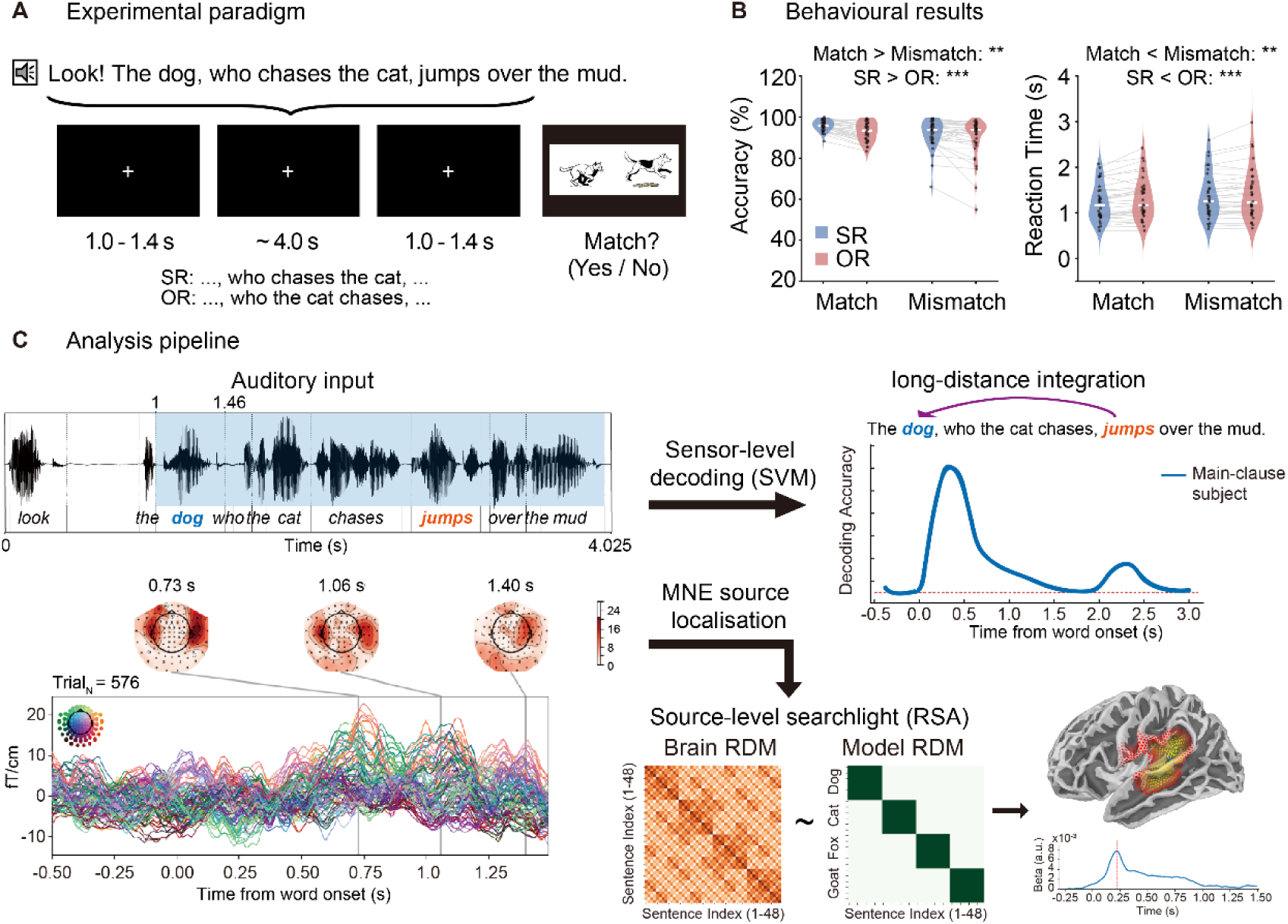
Experimental design, behavioural results, and analysis pipeline. **(A)** Trial structure. Participants listened to spoken sentences with subject-relative (SR) and object-relative (OR) clauses while maintaining fixation (e.g., SR: “The dog, who chases the cat, jumps over the mud”; OR: “The dog, who the cat chases, jumps over the mud”). After each sentence, a probe image appeared and participants judged whether it matched the sentence meaning. **(B)** Behavioural performance. Accuracy (left) and reaction times (right) for Match and Mismatch conditions, shown separately for subject-relative (SR, blue) and object-relative (OR, red) sentences. Dots represent individual participants, and grey lines indicate within-subject comparisons. Both measures showed main effects of congruency and relative-clause type, with higher accuracy and faster responses for Match than Mismatch trials and for SR than OR sentences (two-way repeated-measures ANOVA, **: *p* < 0.01, ***: *p* < 0.001). **(C)** Analysis pipeline. MEG responses to continuous auditory input were time-locked to word onsets to extract subject- and verb-locked epochs. Time-resolved multivariate decoding (SVM) was applied at the sensor level to track the temporal dynamics of main-clause subject representations during online sentence processing. Source activity was reconstructed using minimum-norm estimation, followed by source-level searchlight analysis using representational similarity analysis (RSA) to characterize the spatiotemporal distribution of content-specific representations.

Participants showed high overall accuracy in the behavioural task (mean = 92.6%, SD = 5.2%). A two-way repeated-measures ANOVA revealed main effects of both sentence-picture congruency and syntax on accuracy (Match > Mismatch: *F*_(1,33)_ = 9.28, *p* = 0.005, *η_p_*² = 0.22; SR > OR, *F*_(1,33)_ = 22.4, *p* < 0.001, *η_p_*² = 0.40) and reaction times (Match < Mismatch: *F*_(1,33)_ = 14.1, *p* = 0.001, η_p_² = 0.30; SR < OR, *F*_(1,33)_ = 18.7, *p* < 0.001, *η_p_*² = 0.36). Overall, congruent sentence–picture pairs produced higher accuracy and shorter RTs, and subject-relative sentences were processed more easily than object-relative sentences, consistent with previous psycholinguistic findings ^45–47^.

### WM encoding and reactivation of the main-clause subject

To investigate the encoding and maintenance of main-clause subject, we aligned the MEG epochs to subject onset (“dog”) and trained a Support Vector Machine (SVM) classifier to decode subject identity from sensor-level signals using 12-fold cross-validation (Fig. 1C, upper panel). Decoding accuracy increased significantly above chance following subject onset (∼20 ms), returned to baseline after 1.31 s, and increased again after the onset of the main verb (“jumps”) (Fig. 2A, blue line, cluster 1: 0.02–1.31 s, *p*_cluster_ = 4 × 10^-5^; cluster 2: 2.33–2.64 s, *p*_cluster_ = 0.027; cluster-based permutation test). The decline of decodable information during the embedded clause suggests that the subject representations are maintained in a latent, activity-silent state rather than through sustained activity ^23,24,48^.

**Figure 2.**
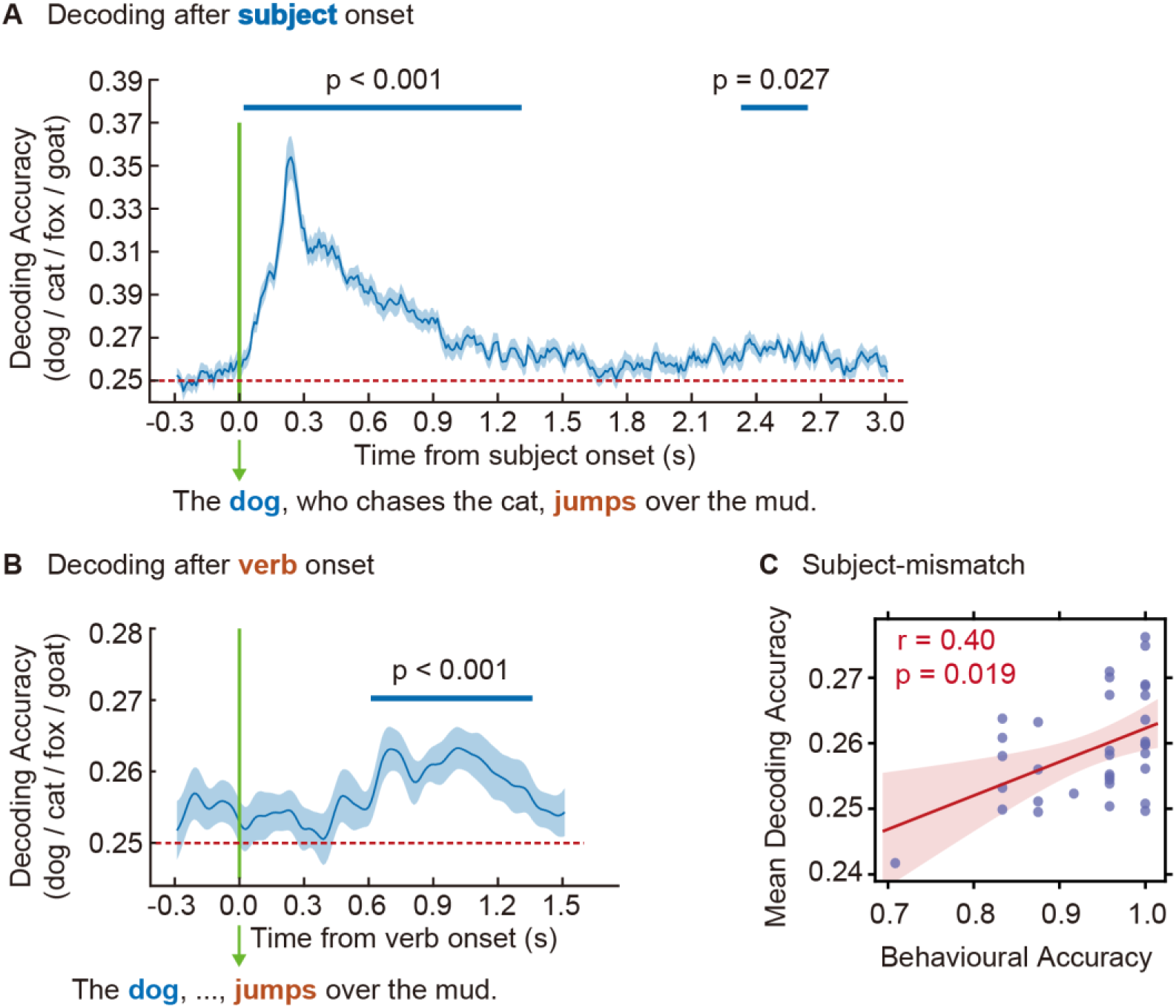
Decoding the main-clause subject across embedded structure. **(A)** Time-resolved decoding aligned to subject onset. Four-way SVM classification of subject identity (dog / cat / fox / goat) revealed significant decoding following subject onset (“dog”) (cluster 1: 0.02–1.31 s, p < 0.001; cluster-based permutation test). A later significant cluster was also observed in the post-verb period (“jumps”) (cluster 2: 2.33–2.64 s, p = 0.027), corresponding to subject reactivation across embedded clauses. Shaded area denotes SEM. Red dashed line indicates chance level (0.25). **(B)** Time-resolved decoding aligned to verb onset. Aligning the epochs to main-verb onset revealed a more precise reactivation window, with subject identity becoming decodable approximately 610 ms after verb onset (“jumps”) (cluster: 0.61–1.36 s, p < 0.001). **(C)** Neural–behavioural correlation. WM reactivation magnitude (mean decoding accuracy 0.5-1.5 s after verb onset) was positively correlated with accuracy on subject-mismatch trials, in which the main-clause subject was replaced by an unmentioned animal (r=0.40, p=0.019, Spearman’s correlation). Each dot represents one participant. Shaded band indicates 95% confidence interval.

To test whether subject reactivation is time-locked to the main verb, we realigned MEG epochs to verb onset (“jumps”). As shown in Fig. 2B, the decoding accuracy of the main subject became significant ∼610 ms after verb onset and remained decodable for approximately 750 ms (Fig. 2B, blue line: 0.61–1.36 s, *p*_cluster_ = 4 × 10^-5^, cluster-based permutation test), demonstrating that the verb triggers memory retrieval of its structure-dependent subject. A neural-behavioural correlation further revealed that reactivation magnitude (mean decoding accuracy 0.5–1.5 s post-verb) was positively correlated with participants’ accuracy in detecting subject-mismatch (Fig. 2C: *r* = 0.40, *p* = 0.019, Spearman’s correlation, N = 34). Together, these findings indicate that long-distance dependency integration relies on the reactivation of subject representations from a latent WM state.

### Representational dynamics underlying WM encoding and reactivation

Our decoding results so far indicate that the main-clause subject is represented during the initial encoding phase and reactivated after verb onset. However, it remains unclear whether the reactivation reinstates the original encoding pattern or involves a transformed representation. To address this, we performed a temporal generalisation analysis ^44^, training SVM classifiers on each time point during encoding (0–1.0 s from subject onset,”dog”) and testing on each time point during reactivation (0–1.0 s from verb onset,”jumps”).

During encoding, we observed a clear diagonal component together with robust off-diagonal generalisation at 0.2 to 0.8 s, revealing that the representational format of the subject remained somewhat stable over half a second (Fig. 3A, left panel; black outline, *p* = 0.001, cluster-based permutation test). In contrast, classifiers trained during subject encoding failed to generalise to the interval reactivated by the verb (Fig. 3A right panel, no significant cluster was found, p > 0.05), and classifiers trained during verb reactivation also failed to generalise to the earlier subject encoding interval (Supplementary Materials Fig. S1). Thus, although subject identity can be decoded during both phases, the lack of cross-phase generalisation implies that reactivation does not reinstate the original encoding pattern but instead recruits a transformed code, reflecting the dynamic nature of language online processing.

**Figure 3.**
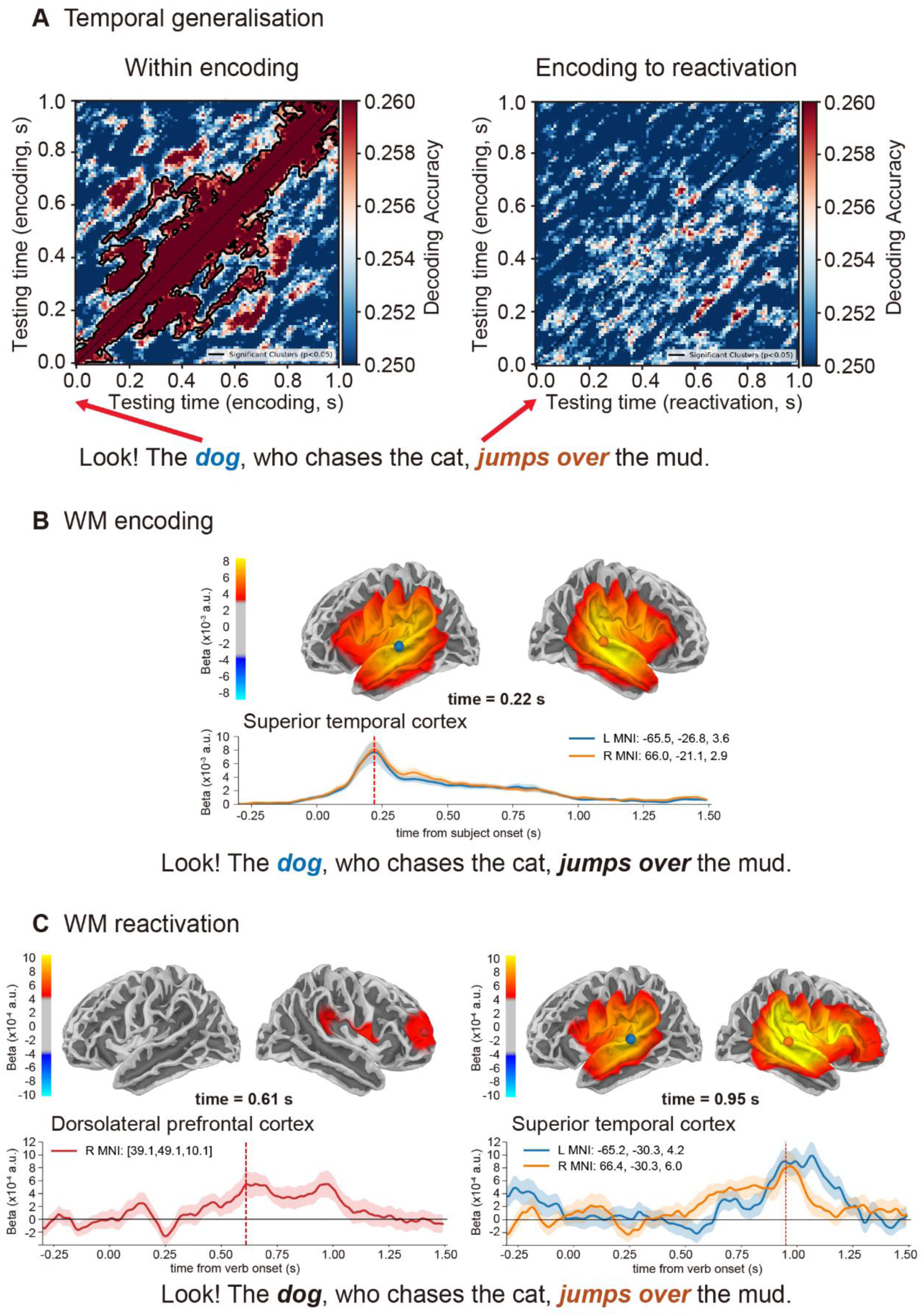
Temporal generalization and source localisation of subject representations. **(A)** Temporal generalisation analysis. Left, classifiers trained and tested within the encoding window (0–1.0 s from subject onset) showed significant within-phase generalisation (black outline, cluster-based permutation test, p = 0.001). Right, classifiers trained during encoding and tested during reactivation (0–1.0 s from verb onset) showed no significant cross-phase generalisation (cluster-based permutation test, p > 0.05). **(B)** Source-level searchlight results during encoding. Cortical map shows the spatial distribution of content-specific effects at 220 ms after subject onset. Time courses are shown for representative sites in left (blue) and right (orange) superior temporal cortex. **(C)** Source-level searchlight results during reactivation. Cortical maps show the spatial distribution of content-specific effects at 610 ms (left panel) and 950 ms (right panel) after verb onset. Time courses are shown for representative sites in the right DLPFC (left panel, red), left superior temporal cortex (right panel, blue), and right superior temporal cortex (orange). In (B) and (C), vertical lines in time-course plots mark the displayed time points and the shaded area denotes SEM.

### Neuronal sources underlying WM encoding and reactivation

We next examined which cortical regions were involved in the WM encoding and reactivation during sentence comprehension. To address this, single-trial MEG data were estimated in source space using minimum-norm estimation (MNE) constrained to the individual cortical surface. Then we applied a searchlight approach to identify the cortical regions that carried content-specific information (for details see Methods).

Source-level searchlight analyses revealed that subject encoding recruited bilateral temporal cortex and inferior frontal gyrus (IFG), with peak decoding accuracy at approximately 220 ms after subject onset (Fig. 3B). During reactivation, content-specific representations initially emerged in right dorsolateral prefrontal cortex (DLPFC), peaking at approximately 610 ms after verb onset (Fig. 3C, left panel), and subsequently engaged temporal cortex and IFG around 950 ms (Fig. 3C, right panel). This temporal progression suggests that verb-triggered WM retrieval is initiated in the prefrontal control network and subsequently propagates to IFG-temporal regions for content representation and subject-verb integration.

### Syntactic structure modulates WM reactivation

Having established the spatiotemporal profile of subject reactivation during subject-verb integration, we next examined whether this process is modulated by the syntactic structure of the embedded clause. MEG epochs were separated into subject-relative (SR) and object-relative (OR) conditions (e.g., SR: “The dog, who chases the cat, jumps over the mud.”, OR: “The dog, who the cat chases, jumps over the mud.”), and SVM decoding analyses were performed separately for each condition.

In the SR condition, subject reactivation emerged 640 ms after verb onset and remained robust for approximately 780 ms (Fig. 4A, left panel: 0.64–1.42 s, *p* = 4 × 10^-5^, cluster-based permutation test). In the OR condition, reactivation occurred later (∼850 ms) and was shorter (∼290 ms) in duration (Fig. 4A, right panel: 0.85–1.14 s, *p* = 0.002, cluster-based permutation test). When comparing the decoding accuracy averaged over the reactivation window identified previously (0.61–1.36 s; see Fig. 2B), we found significantly stronger reactivation in SR than OR sentences (Fig. 4B; paired t-test, *t*_(33)_ = 2.19, *p* = 0.036, Cohen’s *d* = 0.38).

**Figure 4.**
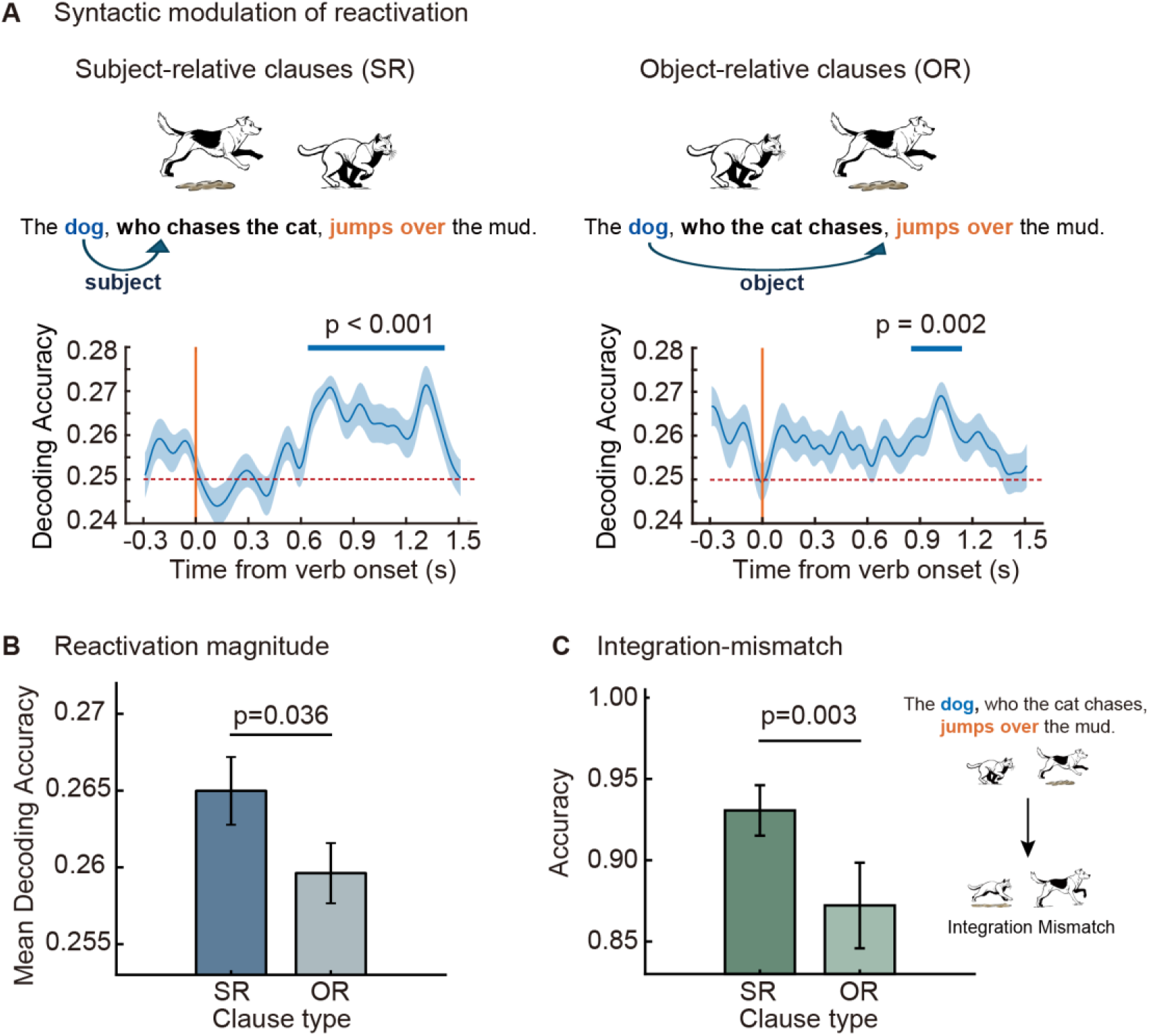
Syntactic structure modulates reactivation and behavioural performance. **(A)** Time-resolved decoding of subject identity for subject-relative (SR, left) and object-relative (OR, right) sentences. Top panels show example sentences with corresponding probe pictures. Blue arrows indicate the syntactic role of the main-clause subject within the embedded clause. Bottom panels show decoding accuracy from −0.3 to 1.5 s relative to verb onset (shaded area denotes SEM). Significant decoding was observed from 0.64-1.42 s in SR sentences (p < 0.001, cluster-based permutation test) and from 0.85-1.14 s in OR sentences (p = 0.002). **(B)** Reactivation magnitude by clause type. Mean decoding accuracy within the reactivation window was higher for SR than OR sentences (paired t-test, p = 0.036). Error bars indicate SEM. **(C)** Accuracy on integration-mismatch trials by clause type. Accuracy was higher for SR than OR sentences (paired t-test, p = 0.003). The right panel illustrates matching and integration-mismatch probe pictures for the example sentence. Error bars indicate SEM.

Behavioural performance showed a similar pattern. A subset of trials was constructed as integration-mismatch trials, in which the probe picture depicted the embedded-clause noun, rather than the main-clause subject, performing the main-clause action, while all other aspects of the sentence remained unchanged. This manipulation selectively violated the correct subject–verb integration (see Fig. 4C for illustration). Accuracy on integration-mismatch trials was higher for SR than OR sentences (Fig. 4C; paired t-test, *t*_(33)_ = 3.21, *p* = 0.003, Cohen’s *d* = 0.55), mirroring the neural reactivation effect. Together, these results indicate that the magnitude and timing of verb-triggered subject reactivation are modulated by the syntactic structure of the embedded clause, with consequences for comprehension accuracy.

## Discussion

In this study, we used time-resolved MEG decoding to track how the neural representation of the main-clause subject is maintained and reactivated across hierarchically embedded structures (e.g.,”The dog, who chases the cat, jumps over the mud. Three key findings emerged: (1) subject (“dog”) representations decayed to baseline across the embedded clause and reactivated at the main-clause verb, consistent with activity-silent maintenance; (2) encoding and reactivation patterns did not cross-generalise, indicating that retrieval reflects a transformed rather than reinstated representation; and (3) reactivation emerged first in right DLPFC before spreading to frontotemporal regions, suggesting domain-general networks contribute to core language functions during hierarchical sentence processing.

How WM supports language comprehension, particularly the processing of long-distance dependencies and syntactic integration, has long been a central question in psycholinguistics and cognitive neuroscience ^34,35,49,50^. The neural mechanisms underlying this capacity provide the foundation for the recursive structure of human language and the generation of unlimited expressions, while also constraining the structural complexity that can be processed, such as the depth of hierarchical embedding ^1,3,4^. Numerous behavioural studies have examined how increased WM load or interference affects language processing difficulty ^47,51–53^, yet the underlying neural mechanisms remain poorly understood. To advance our understanding of the neurobiology of language comprehension, we combined MEG with MVPA to investigate how linguistic information is maintained in WM during sentence processing.

Our results revealed that decoding accuracy for the main-clause subject returned to baseline during processing of embedded clauses and reactivated by the main-clause verb (Fig. 2). This pattern is consistent with the activity-silent theory of WM, in which encoded representations do not persist through sustained neural firing but instead transition into a latent state ^24,48,54,55^. Converging evidence from brain stimulation studies and computational modelling further suggests that information in this latent state is retained through rapid changes in short-term synaptic plasticity ^22,23,56–58^ and periodically consolidated through transient bursts of activity ^59–61^. Critically, this mechanism of activity-silent storage enables more efficient storage of multiple items while minimising interference between maintained representations and incoming inputs, which would be particularly beneficial for sentence processing where multiple constituents must be held in memory as new words are continuously integrated.

Although word representation became activity-silent during maintenance, it remained accessible in WM for later integration. The main-clause verb, which introduces semantic and syntactic constraints, served as a retrieval cue that triggered subject reactivation ^50,62–64^. This cue-driven reactivation is consistent with previous demonstrations using task-relevant cues ^65,66^ and impulse perturbation ^54,56,67^. Critically, WM reactivation in sentence processing was not determined simply by recency or memory trace strength. In subject-relative clauses (e.g.,”The dog, who chases the cat, jumps over the mud.”), the embedded-clause noun (“cat”) was closer to the main-clause verb (“jumps”) and thus constituted a stronger recent competitor ^52^, yet reactivation of the more distant main-clause subject (“dog”) was stronger than in object-relative clauses (e.g.,“The dog, who the cat chases, jumps over the mud.”, Fig. 4). This pattern demonstrated that memory retrieval was guided primarily by syntactic structure rather than by recency ^68^.

Having established structure-dependent WM reactivation at the syntactic integration point, we next asked whether this reactivated representation preserves the same format as the original encoding. Temporal generalisation analysis showed that encoding and reactivation patterns did not cross-generalise (Fig. 3A), indicating that WM retrieval instead engaged a distinct representational state. One plausible interpretation is that encoding and retrieval privilege different aspects of the same linguistic item. During encoding, processing is dominated by word-form information such as phonological features, whereas during retrieval, processing may operate primarily on conceptual information that is relevant for integration with the verb ^64,69^.

Beyond the transformation driven by different cognitive demands at encoding and retrieval, the subject representation may be further reshaped by the intervening context during maintenance. The contrast between sentence structures directly supports this prediction. In subject-relative clauses, the main-clause subject retains its syntactic role throughout, whereas in object-relative clauses it is temporarily reassigned as the object of the embedded verb. Consistent with this structural difference, verbs following subject-relative clauses triggered stronger WM reactivation, a pattern paralleled by higher integration accuracy (Fig. 4). These findings further suggest that the function of WM in sentence comprehension is not merely to preserve prior representations, but to utilise them within a processing memory system that supports their transformation and retrieval during online integration ^5,6^.

A central debate in the cognitive neuroscience of language concerns whether WM operations supporting sentence comprehension are implemented within language-selective regions or rely on domain-general WM networks. Prior work has argued that language WM is primarily located in a left-hemisphere frontotemporal language network (i.e. left inferior frontal gyrus and left superior/middle temporal gyrus), that is functionally distinct from domain-general regions such as DLPFC ^31,33,35^. However, the present findings argue against this dissociation. During long-distance integration, content-specific information was decodable in DLPFC prior to its emergence in classical language regions. This finding result suggests that domain-general WM network contribute directly to core linguistic computation as in non-linguistic WM tasks ^33,70^. The observed right lateralisation suggests that memory retrieval during sentence comprehension extends beyond the left-dominant language-selective network, consistent with evidence that right hemisphere regions support executive functions essential for language processing ^71–74^.

The role of DLPFC in reactivation can be further understood in the context of ongoing debates about its contribution to WM. While early work emphasised DLPFC as a storage site based on sustained activity during maintenance ^18,19^, recent evidence suggests DLPFC primarily supports cognitive control functions such as selection, monitoring, and manipulation of maintained representations ^17,75–77^. Neuroimaging studies distinguishing encoding and retrieval phases have reported stronger DLPFC engagement during retrieval ^78^. Crucially, our findings extend this account by showing that DLPFC carries content-specific information during WM reactivation, indicating that control networks can transiently represent task-relevant content rather than merely operating on representations stored elsewhere.

Taken together, these results suggest that DLPFC contributes to the selection and transformation of representations required for syntactic integration ^12,69^. In this view, reactivation reflects a coordinated process in which domain-general prefrontal regions prioritise and reshape relevant information before it is integrated within language-selective regions ^64^. This pattern aligns with the Memory-Unification-Control framework ^30,49^, which emphasises dynamic interactions between representational and cognitive control systems during sentence comprehension.

Our findings support a model in which WM for hierarchical sentence processing relies on activity-silent maintenance, structure-dependent retrieval, and coordinated engagement of domain-general and language-selective networks. The transformation of word representations across hierarchical structure indicates that WM during language comprehension is not a passive storage buffer but an active system that continuously reshapes representation in accordance with evolving integration demands. These results raise several questions for future investigation. First, we employed controlled sentences structures with fixed embedded types. Future research using more naturalistic materials could test whether similar maintenance and reactivation dynamics generalise across diverse syntactic constructions and embedding depths. Second, the transformation of representations during reactivation bear resemblance to attention mechanisms in transformer-based language models, e.g., GPT-2 ^79,80^. Investigating the alignment between neural reactivation in human brain and attention mechanisms in language models could reveal shared or different principles underlying biological and artificial language processing. Finally, individual differences in reactivation strength predicted comprehension accuracy, raising the possibility that failures of memory retrieval contribute to comprehension difficulties in clinical populations. Examining reactivation dynamics in individuals with language impairments or reduced WM capacity could further identify the role of WM mechanisms in language disorders.

## Acknowledgments

This work was supported by the Wellcome Trust Discovery Award (grant number 227420) and the NIHR Oxford Health Biomedical Research Centre (NIHR203316) to O.J., and the Leverhulme Early Career Fellowship (ECF-294-2023-626) to Y.P. The Oxford University Centre for Integrative Neuroimaging was supported by core funding from the Wellcome Trust (203139/Z/16/Z and 203139/A/16/Z). The views expressed are those of the authors and not necessarily those of the funders. The funders had no role in study design, data collection and the preparation of the manuscript or the decision to publish. We are grateful to Jonathan Winter for assistance with MEG recordings. We thank Sanjay Manohar for his helpful comments.

## Author contributions

J.L. and O.J. conceived and designed the study. Y.P., H.P., and H.L. contributed to the experimental design. J.L. collected and analysed the data. P.H. provided guidance on data analysis and theoretical framing. J.L. and O.J. wrote the manuscript. All authors contributed to interpretation of the results and revision of the manuscript.

## Conflicts of interests

The authors declare no conflicts of interests.

## Data and materials availability

All data and codes will be made publicly available upon acceptance of the paper.

## Materials and methods

### Participants

We recruited 38 native English speakers with normal hearing (screened using the Mimi Hearing Test, or another hearing-test application when Mimi was unavailable) and normal or corrected-to-normal vision. All participants were right-handed and reported no history of neurological problems or diagnosed language disorder. Two participants were discarded because of non-removable dental braces, and two other participants were excluded from analysis because they fell asleep during the MEG session, leaving a final sample of 34 participants (20 females and 14 males, 21.0 ± 3.0 years old, mean ± SD). This sample size was chosen to be comparable to previous studies with a similar experimental design ^26,81,82^. The study was approved by the University of Birmingham Ethics Committee (under the approved Programme ERN_18-0226P). Written informed consent was obtained from all participants after the experimental paradigm and procedures had been explained. Participants received £15 per hour or course credit as compensation for their participation in the MEG and MRI sessions.

### Stimuli and tasks

#### Auditory stimuli (sentences)

In the experiment we constructed 48 unique sentences (see Supplementary Table 1, combining four animals (“dog”,“cat”,“fox”,“goat”) as main-clause subjects with two main-clause verb phrases (“jumps over the mud”,“steps into the mud”)). Besides, the set of natural sentences comprised subject-relative clauses (SR;”who chases …”) and object-relative clauses (OR;”who … chases”), in which the relative clause intervened between the main-clause subject and verb to form a long-distance dependency. The animal within the relative clauses was also chosen from the four options (“dog”,“cat”,“fox”,“goat”) but was never identical to the main-clause subject. This resulted in 48 unique sentences in total (4 main-clause subjects × 2 main-clause verb phrases × 2 relative-clause types × 3 relative-clause nouns).

The experiment was divided into 12 blocks, with each block containing all 48 unique sentences presented in a randomised order, resulting in a total of 576 trials. All auditory stimuli were pre-generated using the text-to-speech service in Vertex AI (Google Cloud, https://console.cloud.google.com/vertex-ai/studio/speech/). We used an English male voice, with sentences presented at approximately 150-160 words per minute (4.0-4.3 s per sentence). Audio files were then converted to MATLAB format for presentation with Psychtoolbox-3, with a sampling rate of 48 kHz. During the MEG session, sounds were delivered via the Windows WASAPI audio subsystem (actual sampling rate: 48 kHz; output latency: 10 ms).

#### Visual stimuli (cartoon illustrations)

After sentence offset on each trial, participants viewed a cartoon illustration on a white background. In half of the blocks (the one-animal blocks), the picture depicted a single animal performing an action (“*jumps over / steps into the mud”*). Then, participants judged whether this event matched or mismatched the main clause of the sentence. In the other half of the blocks (the two-animal blocks), the picture illustrated two animals arranged from left to right in a chasing scene, with one animal also performing the”*jumps over / steps into the mud”* action. Here, participants judged whether the entire event matched or mismatched the full sentence.

Across trials, four possible animals could appear (“*dog”,“cat”,“fox”,“goat”*). Half of the trials were match trials, in which the picture exactly corresponded to the described event, and half were mismatch trials, in which a single aspect of the event was altered. In one-animal blocks, mismatches arose from (1.1) an animal that had not been mentioned in the sentence, (1.2) an action that did not correspond to the sentence, or (1.3) the embedded-clause animal performing the critical action. In two-animal blocks, mismatches arose from (2.1) replacing the main-clause animal with an unmentioned animal, (2.2) replacing the embedded-clause animal with an unmentioned animal, (2.3) swapping the chasing roles of the two animals, or (2.4) assigning the”*jumps over / steps into”* action to the embedded-clause animal rather than the main-clause animal (Supplementary Materials Fig. S2). Only one type of discrepancy was introduced on any given trial. All images used in the experiment were manually drawn.

#### Experimental procedure

Participants were seated in a dimly lit magnetically shielded room under the MEG gantry (60° upright angle) and viewed stimuli on a rear-projection screen positioned 100 cm from their eyes. Visual stimuli were presented by a PROPixx DLP LED projector (VPixx Technologies, Canada), yielding approximately 41 pixels per visual degree. The experimental task was controlled using Psychtoolbox-3 running in MATLAB (MathWorks). Before the main experiment, participants completed a resting-state recording of at least 3 min while maintaining fixation. In addition, an empty-room recording of at least 3 min was acquired on the same day.

Each trial started with a white fixation cross (1.0° of visual angle) at the centre of a black screen (RGB: [0, 0, 0]), lasting for 1.0–1.4 s. An English sentence with an embedded clause (e.g.,“The dog, who chases the cat, jumps over the mud.”) was then presented auditorily while the fixation cross remained on the screen. Auditory stimuli were delivered via a MEGIN/Natus sound system using transducers and air-conduction tubes. After sentence offset, the fixation cross was displayed for another 1.0–1.4 s. Subsequently, a cartoon illustration (31.0°×10.7° visual angle) was presented on a black background (Fig. 1A). Participants then indicated whether the illustration matched the sentence by pressing a button (right index/middle finger). The experiment consisted of twelve blocks, with each block containing 48 sentences and taking approximately 6 min to finish. Following each block, participants were given a rest period of at least 1 min. In total, the MEG recording took about 1.5 h. Thirty-one participants completed all 12 blocks, one participant completed 10 blocks and two participants completed 8 blocks due to early dropout.

### Data acquisition

#### MEG recordings

MEG data were acquired with a 306-sensor TRIUX MEGIN system (204 orthogonal planar gradiometers and 102 magnetometers, Elekta, Finland) installed in a magnetically shielded room. Using a Polhemus Fastrak digitiser (Polhemus Inc., USA), we recorded the location of three fiducial points (nasion, left/right preauricular points), the four HPI coils (two on the mastoids and two on the forehead, separated by at least 3 cm), and more than 200 additional scalp points for later MEG–MRI co-registration. The positions of the HPI coils were recorded in the MEG system and checked at the beginning of each block to ensure that head movement did not exceed 4 mm. The MEG data were online band-pass filtered between 0.1 and 330 Hz to prevent aliasing and sampled at 1,000 Hz.

#### MRI recordings

T1-weighted structural images were acquired using a Siemens 3.0T PRISMA scanner (TR = 2000 ms, TE = 2.01 ms, TI = 880 ms, flip angle = 8°, FOV = 256 × 256 × 208 mm, isotropic voxel size = 1 mm). For two participants who did not undergo MRI scanning, source reconstruction was performed using the FreeSurfer”*fsaverage”* template brain ^83^.

### Behavioural data analysis

#### Accuracy and reaction time

For each participant, trials were assigned to four conditions in a 2 × 2 design crossing Match (match vs mismatch) and Syntax (subject-relative, SR vs object-relative, OR). Mean accuracy across participants (N = 34) was computed in each condition (Fig. 1B, left). Mean reaction time (RT) was computed in the same four conditions, but restricted to correct trials (Fig. 1B, right). Accuracy and RT were analysed separately using two-way repeated-measures ANOVAs, implemented in IBM SPSS Statistics 22 (IBM Corp., Armonk, NY, USA).

In a further analysis, we examined performance in the two-animal blocks when the picture incorrectly bound the animal to the action (error type 2.4), targeting errors of integrating the main-clause subject with its verb (integration-mismatch; Fig. 4C). For each participant, mean accuracy in this integration-mismatch condition was computed separately for SR and OR sentences and compared using a paired-samples t test.

### MEG data analysis

#### Pre-processing

MEG data were pre-processed in Python (version 3.8.2) using MNE-Python (MNE version 1.0.3) ^84^, following the FLUX MEG analysis pipeline ^85^ (https://www.neuosc.com/flux). MaxFilter signal space separation (SSS) was not applied, as previous work has shown that SSS does not necessarily improve sensitivity in multivariate pattern analyses ^86^. The continuous MEG data were firstly down-sampled to 200 Hz and band-pass filtered between 1 and 40 Hz. Independent component analysis (ICA) was subsequently applied ^87^. Then, the ICA components corresponding to eye blinks, saccades, and heartbeat were identified through visual inspection (typically 2 to 4 components per participant) and removed from raw data.

#### Sensor-level decoding analysis

After pre-processed MEG data were epoched relative to sentence onset (marked in the trigger channel), word onsets within each sentence were then obtained by forced-aligning the audio recordings with their orthographic transcripts using Montreal Forced Aligner (MFA, version 3.1.1) with the english_us_arpa acoustic and pronunciation models ^88^. These onset times (relative to sentence onset) were used to extract segments of the MEG epochs aligned to the main-clause subject (Fig. 2A, Fig. 3AB) and to the main verb (Fig. 2B, Fig. 3AC, Fig. 4A), yielding subject-locked and verb-locked data for subsequent decoding analyses. The pre-processed epoch data were low-pass filtered at 20 Hz and then down-sampled to 100 Hz for further analysis.

Time-resolved multivariate pattern classification was then implemented using a support vector machine (SVM; scikit-learn, version 0.23.2) with a radial basis function kernel (RBF) and class weights balanced according to class frequency ^89^. For each time point, feature vectors were constructed by concatenating the signals from all 204 planar gradiometers and 102 magnetometers over a sliding window of 5 consecutive samples (50 ms at 100 Hz), with each time point indexed to the centre of the window, yielding 1,530 features. The decoding pipeline comprised vectorisation and feature-wise z-scoring. The classifier was trained to discriminate the identity of the main-clause subject (four animal categories) using 12-fold stratified cross-validation, providing a time-resolved measure of decoding performance.

Decoding time courses were plotted as mean classification accuracy across participants, with chance at 0.25 (four-way classifier). Statistical testing used a non-parametric sign-permutation test ^54,90^ on accuracy minus chance at each time point (50,000 permutations; two-sided *p* < 0.05). Multiple comparisons over time were controlled with a cluster-based permutation (cluster threshold *p* = 0.05), applied to −0.3–3.0 s around subject onset and −0.3–1.5 s around verb onset. Decoding time courses following verb onset were smoothed using a Gaussian-weighted window (window length = 30 ms) to improve signal-to-noise ratio.

#### Temporal generalisation

To test whether WM encoding and reactivation relied on a shared representational format, we performed a temporal generalisation decoding analysis ^44,54^. We applied the same multivariate decoding approach as in the time-resolved analyses, except that a classifier trained at each encoding time point was tested at all time points across the encoding and reactivation windows, yielding two two-dimensional train-by-test time decoding matrices (Fig. 3A). Diagonal elements of the temporal generalisation matrix correspond to the time-resolved decoding analysis, whereas off-diagonal elements reflect the similarity of content-specific representations across different time points. Statistical significance was assessed at the group level using a cluster-based permutation test on the time × time decoding matrices (one-sample t-tests, one-side *p* < 0.05). The null distribution was obtained by randomly flipping the sign of the decoding accuracy across 10,000 permutations. Clusters with cluster-level *p* < 0.05 were retained.

#### Source reconstruction

Individual T1-weighted structural MRIs were processed with FreeSurfer to reconstruct cortical surfaces and to compute a boundary element model (BEM) using the watershed algorithm ^91^. MEG sensor locations were co-registered to each participant’s MRI, and a cortical constrained source space was defined on the reconstructed surfaces. Forward solutions were computed including all planar gradiometers and magnetometers. The noise covariance matrix was estimated from 30 s of empty-room recordings acquired on the experimental day, using a shrinkage estimator and subsequent regularisation for magnetometers and gradiometers (mag = 0.2, grad = 0.2; rank based on the MEG info). This covariance matrix was used to spatially pre-whiten magnetometers and gradiometers prior to inverse modelling, thereby accounting for differences in sensor scaling and noise levels. A minimum-norm inverse operator was then constructed in MNE-Python, with source orientations constrained to be normal to the cortical surface. This inverse operator was applied to single epochs to obtain dynamic statistical parametric maps (dSPM source estimates) for each trial ^92^. A surface-based source space consisting of 8196 vertices (4098 per hemisphere) was generated for each participant (Fig. 1C).

#### Source-level searchlight analysis

We performed whole brain searchlight using representational similarity analysis (RSA) ^93^ on the single-trial source estimates with the MNE-RSA toolbox (mne_rsa version 1.0) ^94^. RSA was chosen over SVM-based decoding because RSA is more robust in high-dimensional, low-sample-size settings typical of neuroimaging searchlight analyses, and is computationally more efficient when testing multiple searchlight spheres across the brain. For each participant, we first constructed a model representational dissimilarity matrix (RDM) that captured categorical differences between main-clause subjects: for every pair of trials, entries were coded as 0 when the main-clause subject category was the same and 1 when it differed. Around each source location, a searchlight patch was defined with a spatial radius of 5 cm and a temporal radius of 50 ms, resulting in 1639200 searchlight patches. Within each spatio-temporal patch, a neural RDM was constructed using squared Euclidean distance between trial-wise source patterns. This neural RDM was then correlated with the model RDM using Kendall’s tau-a, yielding a time-resolved regression coefficient value for each source location within a predefined analysis interval (–0.3 to 1.5 s) for analyses time-locked to subject and verb onset, respectively (see Fig. 3BC).

## Supplementary materials

**Supplementary Table 1.** Set of 48 experimental sentences and indexes.

| Sentence ID | Sentence | Main-clause subject (index) | Relative-clause noun (index) | Syntax (index) | Verb phrase (index) |
| --- | --- | --- | --- | --- | --- |
| 1 | Look! The cat, who the dog chases, jumps over the mud. | 1 | 2 | 2 | 1 |
| 2 | Look! The cat, who chases the dog, jumps over the mud. | 1 | 2 | 1 | 1 |
| 3 | Look! The cat, who the dog chases, steps into the mud. | 1 | 2 | 2 | 2 |
| 4 | Look! The cat, who chases the dog, steps into the mud. | 1 | 2 | 1 | 2 |
| 5 | Look! The cat, who the fox chases, jumps over the mud. | 1 | 3 | 2 | 1 |
| 6 | Look! The cat, who chases the fox, jumps over the mud. | 1 | 3 | 1 | 1 |
| 7 | Look! The cat, who the fox chases, steps into the mud. | 1 | 3 | 2 | 2 |
| 8 | Look! The cat, who chases the fox, steps into the mud. | 1 | 3 | 1 | 2 |
| 9 | Look! The cat, who the goat chases, jumps over the mud. | 1 | 4 | 2 | 1 |
| 10 | Look! The cat, who chases the goat, jumps over the mud. | 1 | 4 | 1 | 1 |
| 11 | Look! The cat, who the goat chases, steps into the mud. | 1 | 4 | 2 | 2 |
| 12 | Look! The cat, who chases the goat, steps into the mud. | 1 | 4 | 1 | 2 |
| 13 | Look! The dog, who the cat chases, jumps over the mud. | 2 | 1 | 2 | 1 |
| 14 | Look! The dog, who chases the cat, jumps over the mud. | 2 | 1 | 1 | 1 |
| 15 | Look! The dog, who the cat chases, steps into the mud. | 2 | 1 | 2 | 2 |
| 16 | Look! The dog, who chases the cat, steps into the mud. | 2 | 1 | 1 | 2 |
| 17 | Look! The dog, who the fox chases, jumps over the mud. | 2 | 3 | 2 | 1 |
| 18 | Look! The dog, who chases the fox, jumps over the mud. | 2 | 3 | 1 | 1 |
| 19 | Look! The dog, who the fox chases, steps into the mud. | 2 | 3 | 2 | 2 |
| 20 | Look! The dog, who chases the fox, steps into the mud. | 2 | 3 | 1 | 2 |
| 21 | Look! The dog, who the goat chases, jumps over the mud. | 2 | 4 | 2 | 1 |
| 22 | Look! The dog, who chases the goat, jumps over the mud. | 2 | 4 | 1 | 1 |
| 23 | Look! The dog, who the goat chases, steps into the mud. | 2 | 4 | 2 | 2 |
| 24 | Look! The dog, who chases the goat, steps into the mud. | 2 | 4 | 1 | 2 |
| 25 | Look! The fox, who the cat chases, jumps over the mud. | 3 | 1 | 2 | 1 |
| 26 | Look! The fox, who chases the cat, jumps over the mud. | 3 | 1 | 1 | 1 |
| 27 | Look! The fox, who the cat chases, steps into the mud. | 3 | 1 | 2 | 2 |
| 28 | Look! The fox, who chases the cat, steps into the mud. | 3 | 1 | 1 | 2 |
| 29 | Look! The fox, who the dog chases, jumps over the mud. | 3 | 2 | 2 | 1 |
| 30 | Look! The fox, who chases the dog, jumps over the mud. | 3 | 2 | 1 | 1 |
| 31 | Look! The fox, who the dog chases, steps into the mud. | 3 | 2 | 2 | 2 |
| 32 | Look! The fox, who chases the dog, steps into the mud. | 3 | 2 | 1 | 2 |
| 33 | Look! The fox, who the goat chases, jumps over the mud. | 3 | 4 | 2 | 1 |
| 34 | Look! The fox, who chases the goat, jumps over the mud. | 3 | 4 | 1 | 1 |
| 35 | Look! The fox, who the goat chases, steps into the mud. | 3 | 4 | 2 | 2 |
| 36 | Look! The fox, who chases the goat, steps into the mud. | 3 | 4 | 1 | 2 |
| 37 | Look! The goat, who the cat chases, jumps over the mud. | 4 | 1 | 2 | 1 |
| 38 | Look! The goat, who chases the cat, jumps over the mud. | 4 | 1 | 1 | 1 |
| 39 | Look! The goat, who the cat chases, steps into the mud. | 4 | 1 | 2 | 2 |
| 40 | Look! The goat, who chases the cat, steps into the mud. | 4 | 1 | 1 | 2 |
| 41 | Look! The goat, who the dog chases, jumps over the mud. | 4 | 2 | 2 | 1 |
| 42 | Look! The goat, who chases the dog, jumps over the mud. | 4 | 2 | 1 | 1 |
| 43 | Look! The goat, who the dog chases, steps into the mud. | 4 | 2 | 2 | 2 |
| 44 | Look! The goat, who chases the dog, steps into the mud. | 4 | 2 | 1 | 2 |
| 45 | Look! The goat, who the fox chases, jumps over the mud. | 4 | 3 | 2 | 1 |
| 46 | Look! The goat, who chases the fox, jumps over the mud. | 4 | 3 | 1 | 1 |
| 47 | Look! The goat, who the fox chases, steps into the mud. | 4 | 3 | 2 | 2 |
| 48 | Look! The goat, who chases the fox, steps into the mud. | 4 | 3 | 1 | 2 |

**Figure S1.**
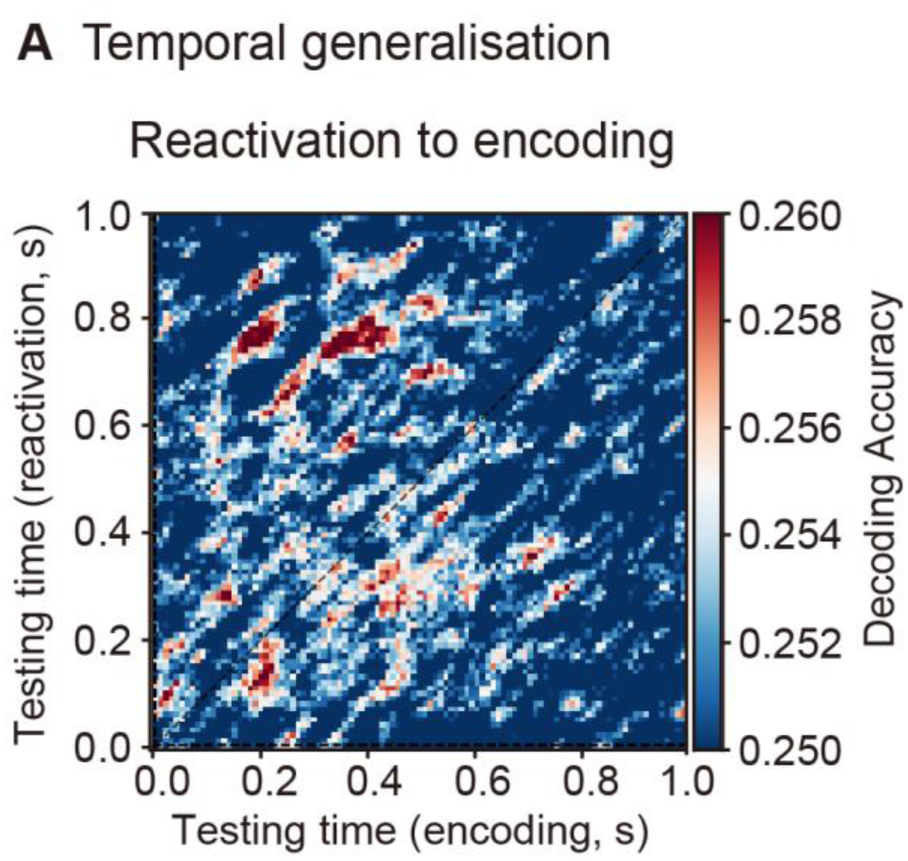
Temporal generalization from reactivation to encoding. **(A)** Temporal generalisation analysis. Classifiers trained during subject reactivation (0–1.0 s from verb onset) and tested during encoding (0–1.0 s from subject onset) showed no significant cross-phase generalisation (cluster-based permutation test, p > 0.05).

**Figure S2.**
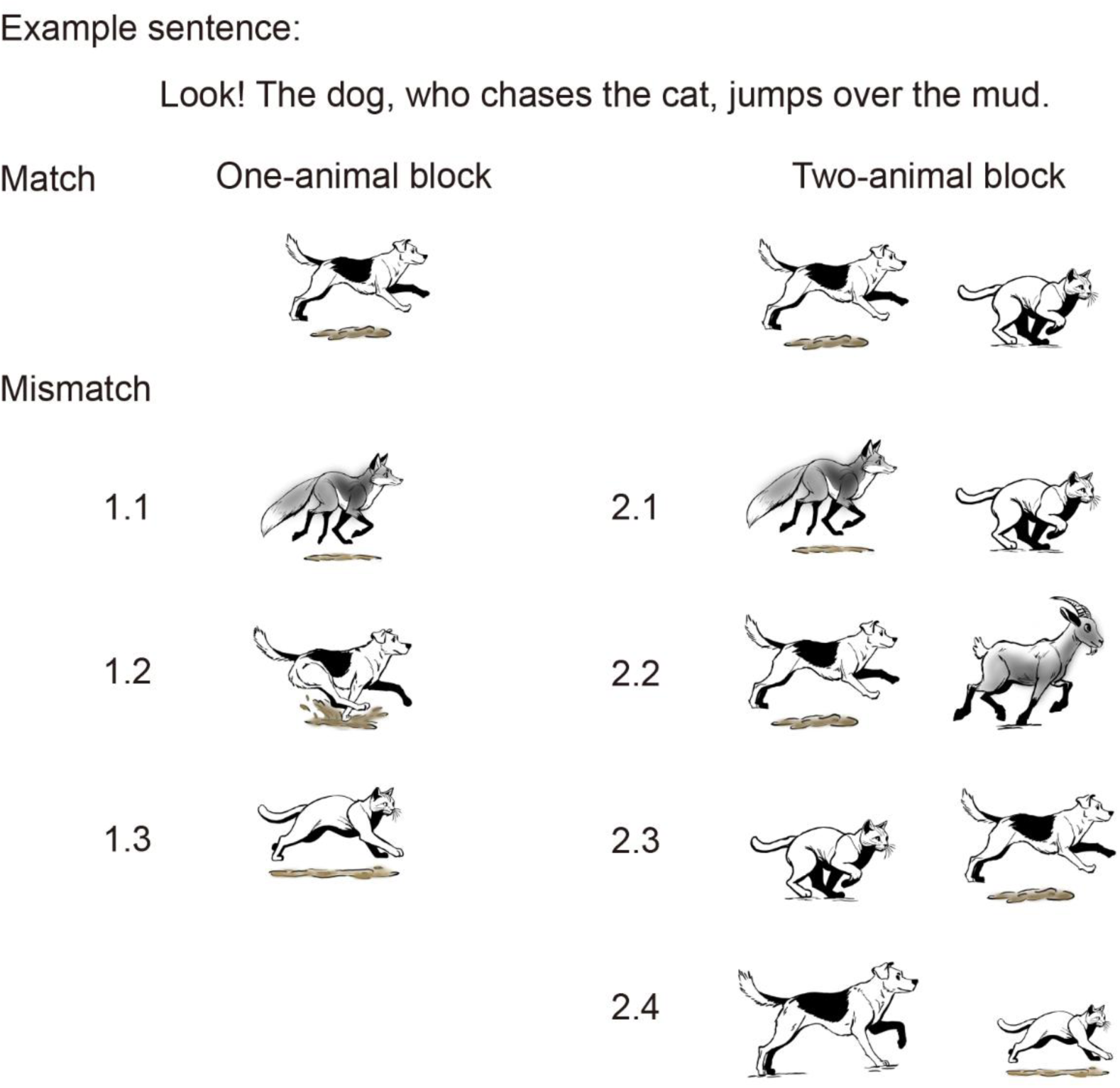
Illustration of stimulus conditions in the sentence-picture matching task. Example sentence and corresponding probe images used in the experiment. Trials were divided into one-animal and two-animal blocks. In match trials, the picture corresponded exactly to the described event. In mismatch trials, a single aspect of the event was altered. In one-animal blocks, mismatches included (1.1) an unmentioned animal performing the action, (1.2) an incorrect action, and (1.3) the embedded-clause animal performing the main-clause action. In two-animal blocks, mismatches included (2.1) replacing the main-clause animal, (2.2) replacing the embedded-clause animal, (2.3) swapping the chasing roles of the two animals, and (2.4) assigning the main-clause action to the embedded-clause animal.

## Notes

### Competing Interest Statement

The authors have declared no competing interest.

